# Managing the Spatial Covariance of Genetic Diversity in Niemann-Pick C1 Through Modulation of the Hsp70 Chaperone System

**DOI:** 10.1101/437764

**Authors:** Chao Wang, Samantha M Scott, Darren M Hutt, Pei Zhao, Hao Shao, Jason E Gestwicki, Balch William E

**Affiliations:** Department of Molecular Medicine and The Skaggs Institute for Chemical Biology, The Scripps Research Institute (Scripps Research), La Jolla; Pharmaceutical Chemistry, University of California, San Francisco, San Francisco, CA

## Abstract

Genetic diversity provides a rich repository for understanding the role of proteostasis in the management of the protein fold to allow biology to evolve through variation in the population and in response to the environment. Failure in proteostasis can trigger multiple disease states affecting both human health and lifespan. Niemann-Pick C (NPC) disease is a genetic disorder mainly caused by mutations in NPC1, a multi-spanning transmembrane protein that is trafficked through the exocytic pathway to late endosomes and lysosomes (LE/Ly) to manage cholesterol homeostasis. Proteostatic defects triggered by >600 NPC1 variants found in the human population inhibit export of NPC1 protein from ER or function in downstream LE/Ly, leading to accumulation of cholesterol and rapid onset neurodegeneration in childhood for most patients. We now show that chemical allosteric inhibitors, such as JG98, targeting the cytosolic Hsp70 chaperone/co-chaperone complex improves the trafficking and stability of NPC1 variants with diverse NPC1 genotypes. By exploiting the knowledge-base of NPC1 variants found in the world-wide patient population using Variation Spatial Profiling (VSP), a Gaussian-process based machine learning (ML) approach, we show how the Hsp70 chaperone system alters the spatial covariance (SCV) tolerance of the ER and the SCV set-points for each residue of the NPC1 polypeptide chain differentially to improve trafficking efficiency and post-ER stability for variants distributed across the entire NPC1 polypeptide. The impact of JG98 is supported by the observation that silencing of Hsp70 specific nucleotide exchange factors (NEF) (BCL-anthogene (BAG) family) co-chaperones significantly improve the folding status of NPC1 variants. Together, these studies suggest that targeting the cytosolic Hsp70 system to adjust the SCV tolerance of the proteostasis network can improve recognition of the plasticity of the NPC1 fold found in the disease population for trafficking to the LE/Ly compartments.

## INTRODUCTION

Understanding how genetic diversity contributes to health and disease in the human population is a problem that continues to confounds medical practice [1, 2]. This challenge has led to the need for a high definition [3] or precision medicine approach (https://allofus.nih.gov/) to address the more than ~10,000 familial and somatic rare diseases [4,] as well as the somatic diseases leading to cancer [5, 6] and the accumulation of variants over a lifespan that lead to numerous neurodegenerative disorders [7, 8].

Niemann-Pick type C (NPC) is an inherited autosomal recessive disorder that is characterized by the accumulation of cholesterol and other lipids in the late endosome (LE) and lysosome (Ly) compartments (LE/Ly) [9–11]. It is caused by mutations in NPC1 or NPC2, with former representing 95% of recorded cases. These two proteins are essential for managing the distribution of cholesterol at the plasma membrane through the LE/Ly cholesterol recycling center. Currently, there are > 600 NPC1 variants recorded in the clinic (https://medgen.medizin.uni-tuebingen.de/NPC-db2/) that result in the accumulation of cholesterol in LE/Ly. Because of the importance of cholesterol homeostasis in the brain [12], this cholesterol accumulation ultimately leads to lethal neurological decline before the age of 25. Currently, there is no cure for NPC disease.

NPC1 is a late endosomal (LE) multi-spanning transmembrane protein responsible for cholesterol efflux out of the lysosome. NPC1 contains three luminal domains (SNLD1, MLD3, CLD5) and three transmembrane domains (NTMD2, STMD4, CTMD6) with 13 helices (Fig 1A)[13]. NPC1 utilizes the middle luminal domain (MLD3) to mediate the transfer of NPC2-bound cholesterol to the sterol binding domain (SNLD1) located at the N-terminal domain of NPC1 [14, 15]. The most common disease-causing variant, I1061T, is located in the CLD5 domain exhibits similar mRNA levels to that seen with wild-type (WT) NPC1, but produces a misfolded protein that is accumulated in and targeted for degradation in the ER, resulting in a >85% deficiency of LE/Ly-localized NPC1 in all tissues, thereby establishing a loss-of-function condition [16]. A compensatory overexpression of NPC1-I1061T, in transgenic NPC1 mouse and cat models, was able to provide a partial correction of the disease phenotype [16]. In agreement with these data, it has also been shown that pharmacological treatments that lead to upregulation of NPC1 variant expression, including ryanodine receptor antagonists [17], treatment with oxysterols that bind to and stabilize NPC1 [18], reduced expression of TMEM97 [19], and HDAC inhibitors [20], partially abrogate the NPC1-I1061T phenotype in heterologous cell and patient fibroblast models. These findings highlight that modulation of the local folding and signaling environment can facilitate disease correction [21–24].

**Figure 1.**
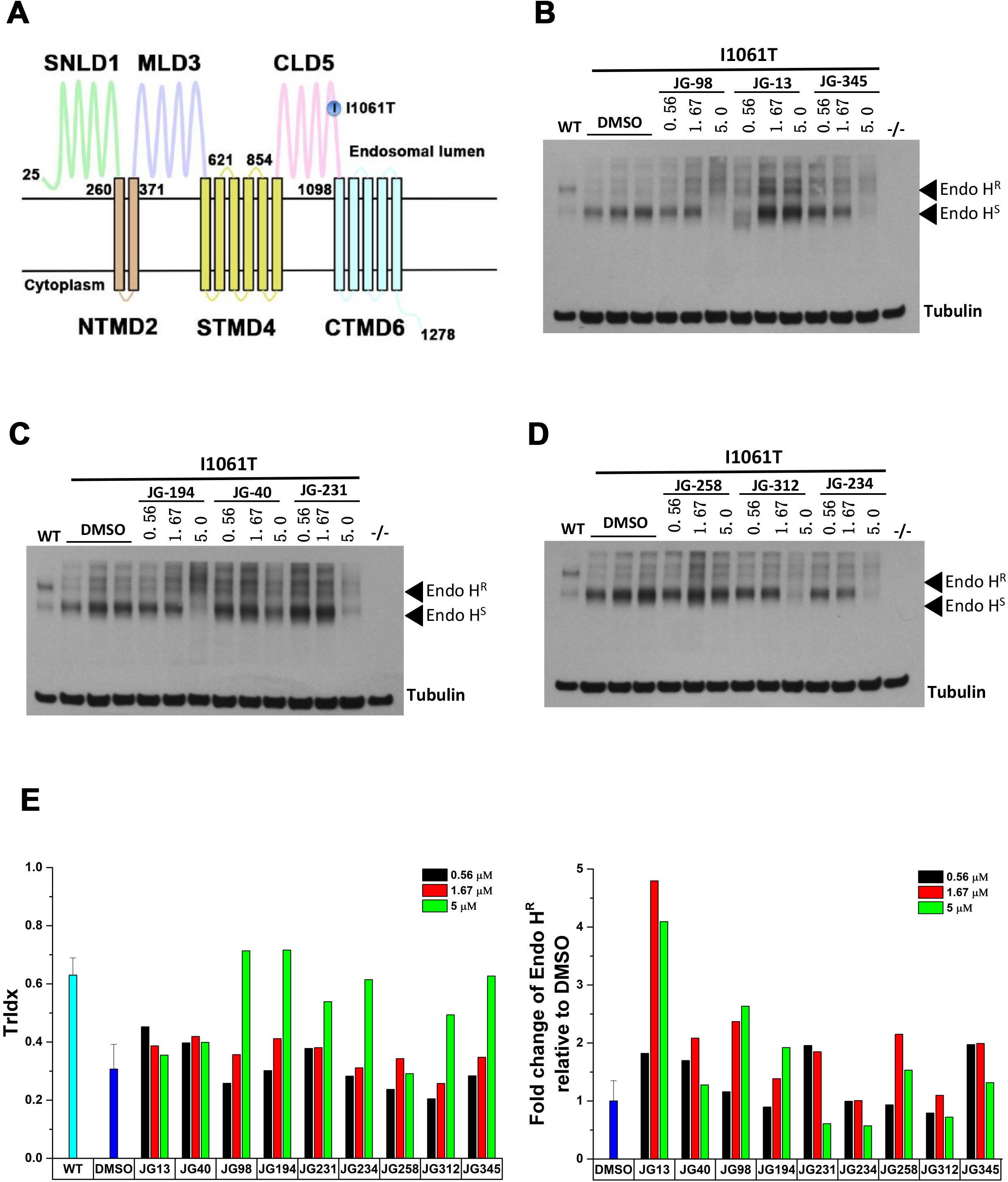
The impact of Hsp70 allosteric inhibitors on the trafficking and stability of NPC1-I1061T. (**A**) Domains in the NPC1 protein. Sterol-binding luminal domain1 (SNLD1), N-terminal transmembrane domain 2 (TMD2), middle luminal domain 3 (MLD3), sterol-sensing transmembrane domain 4 (STMD4), C-terminal luminal domain 5 (CLD5), C-terminal transmembrane domain 6 (CTMD6). (**B-D**) A series of Hsp70 allosteric inhibitors were tested in U2OS-SRA-shNPC1 cells transfected with NPC1-I1061T. All sample lysates were treated with EndoH and analyzed by SDS-PAGE. The slower migrating band is the EndoH^R^ species representative of the post-ER glycoform of NPC1 protein. The faster migrating band is EndoH^S^ species representative of the immature glycoform of NPC1 protein located in the ER. (**E**) Quantification of Western-blot results in B-D. Trafficking index (TrIdx) (left panel) is calculated as EndoH^R^/(EndoH^R^ + EndoH^S^) defining the trafficking efficiency of NPC1-I1061T. Fold-change of EndoH^R^ relative to DMSO after JG98 treatment (right panel) represents the change of protein level of the post-ER glycoforms of NPC1 after compound treatment reported as a specific activity of the total μg protein loaded on the gel.

Proteostasis [8] is responsible for maintaining a healthy proteome environment [7, 25–27] under normal homeostatic conditions and in response to folding stress in each cell type [28]. Protein folding is actively managed, in part, by heat shock cognate (Hsc) and inducible heat shock protein (Hsp) 70 chaperone/co-cochaperone system (referred to collectively as Hsp70) that comprises nearly 5% of cytoplasmic protein pool [16]. The Hsp70 system comprises a network of proteins mediating nascent protein synthesis, folding, stability, degradation, and mediates aggregation [6, 7, 29–31]. Hsp70 is assisted by Hsp40 co-chaperones that stimulate the ATPase activity of Hsp70 [32–36] and nuclear exchange factors (NEFs), such as the BCL-anthogene (BAG) family members, which promote ADP and client protein release [7, 37–39]. The Hsp70 system has been recently suggested to impact the stability of NPC1-I1061T [40–43], but it is not clear how this link might be targeted with small molecules to rescue NPC1 processing. One possibility is a class of allosteric Hsp70 inhibitors, which disrupt the protein-protein interactions between Hsp70 and BAG family co-chaperones [5, 38, 44–47]. These allosteric inhibitors have been shown to promote normal proteostasis in multiple disease models, for example, Dengue virus [48], hepatitis C virus [49], tauopathy [50] and cancer [5, 45, 46].

Herein, we screened a panel of Hsp70 allosteric inhibitors to assess their impact on NPC1-I1061T trafficking. We identified JG98 [6, 29, 38, 45, 48, 49, 51, 52], as a potent stabilizer and trafficking corrector of NPC1-1061T. We also observed that JG98 corrected the trafficking defect seen with 58 additional NPC1 variants, suggesting a more universal but variable impact of this small molecule on NPC1 variants. To understand how Hsp70 manages NPC1 variants distributed across the entire polypeptide chain, we applied Variation Spatial Profiling (VSP) [2], a Gaussian process (GP)-based machine learning (ML) algorithm that utilizes a sparse collection of variants recorded in the world-wide patient population to predict sequence-to-function-to-structure relationships across the entire gene. VSP establishes the phenotypic impact of each amino acid on polypeptide function as a matrix, a biological principle referred to as spatial covariance (SCV) [2]. This approach allows us to establish SCV tolerances of a given protein fold to maintain biological function and assigns SCV set-points to each residue in the polypeptide chain to determine its role in sequence-to-function-to-structure design [2]. Using VSP, we can assign the differential SCV response for trafficking and post-ER stability of the entire NPC1 polypeptide to JG98 [2]. These SCV relationships reveal that JG98 improves recognition by the Hsp70 system of both the modularity and plasticity of the NPC1 fold. These results are supported by our observations showing correction of the trafficking defect of the I1061T variant in response to the silencing of BAGs 1-3, suggesting that SCV set-points assigned to NPC1 residues, in response to the dynamic composition of the Hsp70-NEF-client complex, provide an approach based on the computational formalism of VSP [2] that can leverage the proteostasis program to restore function.

## RESULTS

### Hsp70 allosteric inhibitors improve trafficking and post-ER stability of NPC1-I1016T

NPC1 is a multi-spanning transmembrane protein (Fig 1A) with a prominent luminal orientation dedicated to the management of cholesterol in the LE/Ly compartment [16, 53, 54]. The trafficking of NPC1 can be monitored by assessing the sensitivity or resistance of its 14 N-linked glycans to endoglycosidase H (EndoH) digestion. ER-localized NPC1 glycans are sensitive to EndoH digestion (EndoH^S^) whereas the glycans on NPC1 molecules that are exported to the Golgi are modified by Golgi-resident enzymes and become resistant to EndoH digestion (EndoH^R^) thereby allowing for distinct migration of the retained and exported fractions on SDS-PAGE. ~70% of WT-NPC1 migrates as EndoH^R^ species (Fig 1B), reflecting its steady-state distribution in post-ER compartments. In contrast, more than 80% of NPC1-I1061T exhibits EndoH sensitivity, demonstrating that this misfolded variant is retained in the ER. Here, we quantify these defects in two ways. First, while some band laddering is observed, reflecting the variable processing of the 14-N-linked glycans, we use the EndoH^R^ band seen in WT NPC1 (Fig 1B) as a quantitative measure of ER export for calculating a trafficking index (TrIdx) (EndoH^R^/(EndoH^R^ + EndoH^S^)), a metric which reflects the efficiency of ER export. Second, we quantify the total fold change in EndoH^R^ band intensity as the amount of EndoH^R^ per mg protein compared to a solvent or mock control. This approach reports on the total increase in NPC1 that is transported to the post-ER compartments.

Using this approach, we tested a panel of Hsp70 allosteric inhibitors [5] for effects on the trafficking and stability of NPC1-I1061T. A panel of related molecules was screened because previous studies have shown that the optimal molecule can be subtly different in different cell lines. Accordingly, we tested JG-13, JG-40, JG-98, JG-194, JG-231, JG-234, JG-312 and JG-345, as well as a negative control compound JG258, for effects on TrIdx and EndoH^R^ glycoforms (Fig 1B-D). Quantitation of the TrIdx (Fig 1E, left panel) revealed that 5 μM of JG98 and JG194 lead to a striking increase for NPC1-I1061T, resulting in values that are equivalent to or greater than that observed for WT NPC1. We also observed that JG13 and JG98 at 5 μM markedly improved the fold-change in EndoH^R^ glycoforms, relative to DMSO control (Fig 1E, right panel). Considering both the improvement of TrIdx and EndoH^R^, we decided to use JG98 in further experiments.

### Impact of JG98 on NPC1 variants in patient fibroblasts

To validate the corrective properties of JG98 in a physiologically relevant cellular environment, we monitored the impact of JG98 (5 μM) on patient-derived fibroblasts harboring different NPC1 genotypes (Fig 2A). Of the 18 genotypes tested, 10 exhibited an increased TrIdx in response to JG98, with 7 of these also exhibiting an increase in EndoH^R^ of at least 2-fold and 2 showing increases in EndoH^R^that were less than 2-fold (Fig. 2A-C). In addition to the 7 variants described above, there were 4 other genotypes that exhibited a > 2-fold increase in EndoH^R^ in response to JG98 without a significant increase in their respective TrIdx, namely P401T/I1061T, G673V/I1061T, T1036M/I1061T and I1061T/I1061T (Fig. 2A-C), suggesting that for these genotypes, the primary impact of JG98 is to stabilize the overall level of NPC1 variants. Overall, both TrIdx values (Fig 2B, right panel) and the EndoH^R^ levels (Fig 2C, right panel) of most of the variant population are improved by addition of JG98, although the effects on appearance of EndoH^R^ glycoforms are more pronounced. This result indicates for these fibroblasts, JG98 can increase the NPC1 protein level even if the effects on trafficking are less clear. Consistent with this interpretation, an examination of the response of the I1061T homozygous fibroblast to JG98 (Fig 2D), revealed a dose-dependent increase in the EndoH^R^ (Fig 2E, right panel) which was not observed for the TrIdx (Fig 2E, left panel). Considering that all genotypes tested, except one, were heterozygote and that many exhibited significant responses to JG98, these data suggest that modulation of the Hsp70 chaperone system has a substantial impact on the stability of numerous different NPC1 folding intermediates contributing to an increase in trafficking. How these different NPC1 variants respond individually to JG98-mediated modulation of the Hsp70 machinery will provide insight into the contribution of this chaperone on the biogenesis, stability and trafficking of NPC1.

**Figure 2.**
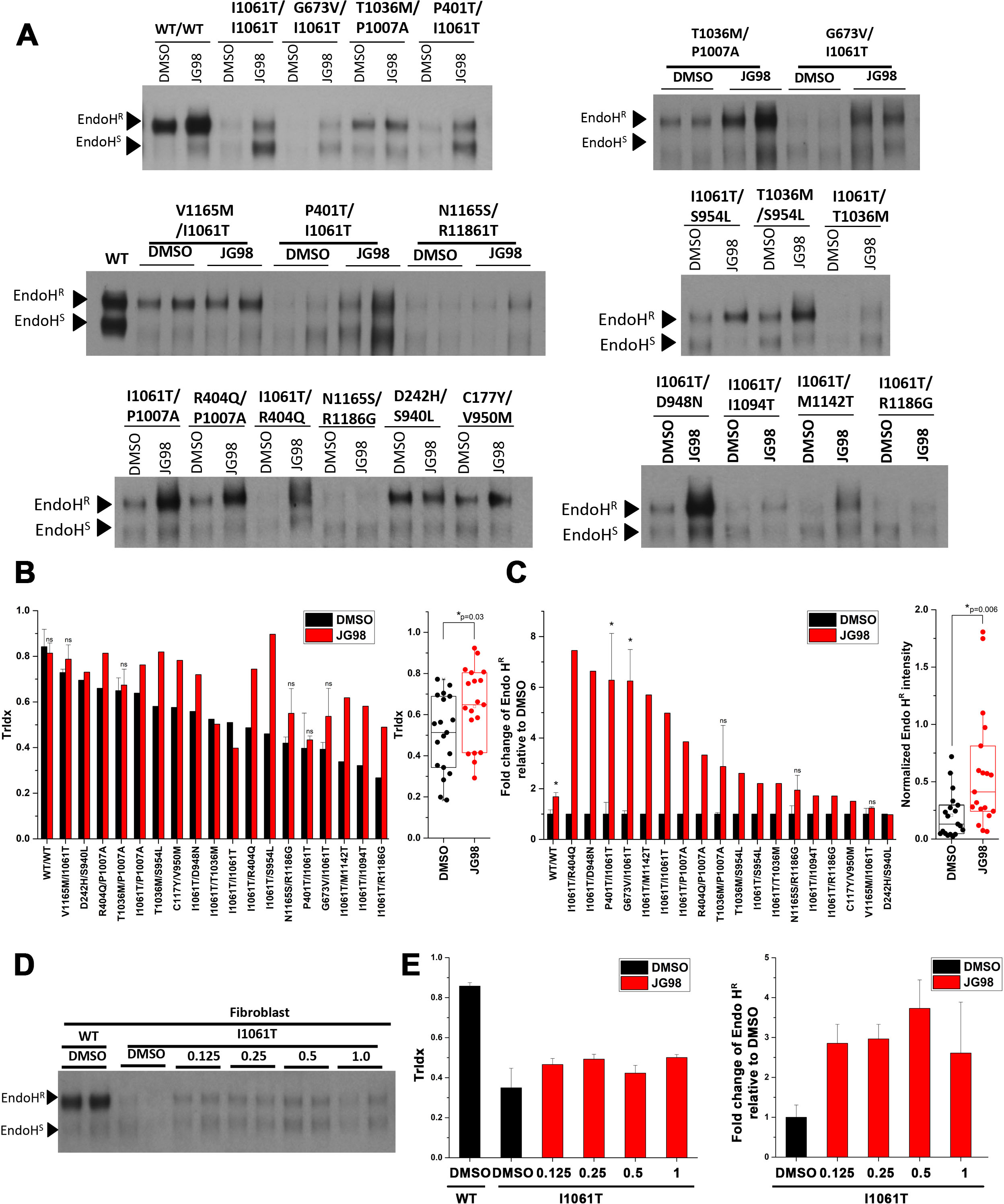
Impact of JG98 on NPC patient fibroblasts. (**A**) JG98 was tested on a collection of fibroblasts from patients with different NPC1 genotypes (see **Methods**). NPC1 was immunoprecipitated from 75 μg of total cell lysate protein, treated with EndoH, eluted from anti-NPC1 beads and separated on a 4-12% Bis-Tris SDS-PAGE. (**B**)Quantification of TrIdx (left panel) for each fibroblast. Four fibroblasts were tested in triplicates and 1 fibroblast<0.05,*; p>0.05, ns; Student’s t-test) are indicated. The remaining 14 fibroblasts are reported as singlets. Fibroblasts with different NPC1 genotypes are ordered according to their TrIdx value in DMSO condition. Right panel shows a box and whisker plot for the TrIdx of all the fibroblasts in the DMSO and JG98 condition. p-value (Student’s t-test) is indicated. (**C**) Quantification of the fold-change of EndoH^R^ species relative to DMSO (left panel). Fibroblasts with different genotypes are arranged according to the fold-change in EndoH^R^ species relative to DMSO. For the fibroblasts that have experimental replicates, error bar (mean ± SD) and p-value (p<0.05,*; p>0.05, ns; Student’s t-test) are indicated. Right panel shows a box and whisker plot for the normalized EndoH^R^ species relative to the WT standard run in each gel. p-value (Student’s t-test) is indicated. (**D**) Analysis of impact of 0.125 μM, 0.25 μM, 0.5 μM, and 1.0 μM JG98 for 24 h fibroblast I1061T/I1061T genotype in duplicate. (**E**) Quantification of TrIdx (left panel) and the fold change of EndoH^R^ relative to DMSO with JG98 treatment (right panel) for the dose experiment shown in (**D**) (mean ± SD).

### Impact of JG98 on NPC1 variants

Patient fibroblasts are mostly heterozygous and have diverse genetic backgrounds and natural history’s given their collection from patients around the world. The complex genetic background make it difficult to dissect the impact of JG98 on the trafficking of individual NPC1 variants. Therefore, we constructed 58 different patient-derived NPC1 variants spanning the entire polypeptide chain using NPC1 ‘WT-V’ sequence [42, 55–59] (Fig 3A). These variants were transiently transfected into NPC1-null U2OS cells [20] for characterization of their baseline, and their JG98-responsive TrIdx (Fig 3B) and the EndoH^R^ glycoform (Fig 3C).

**Figure 3.**
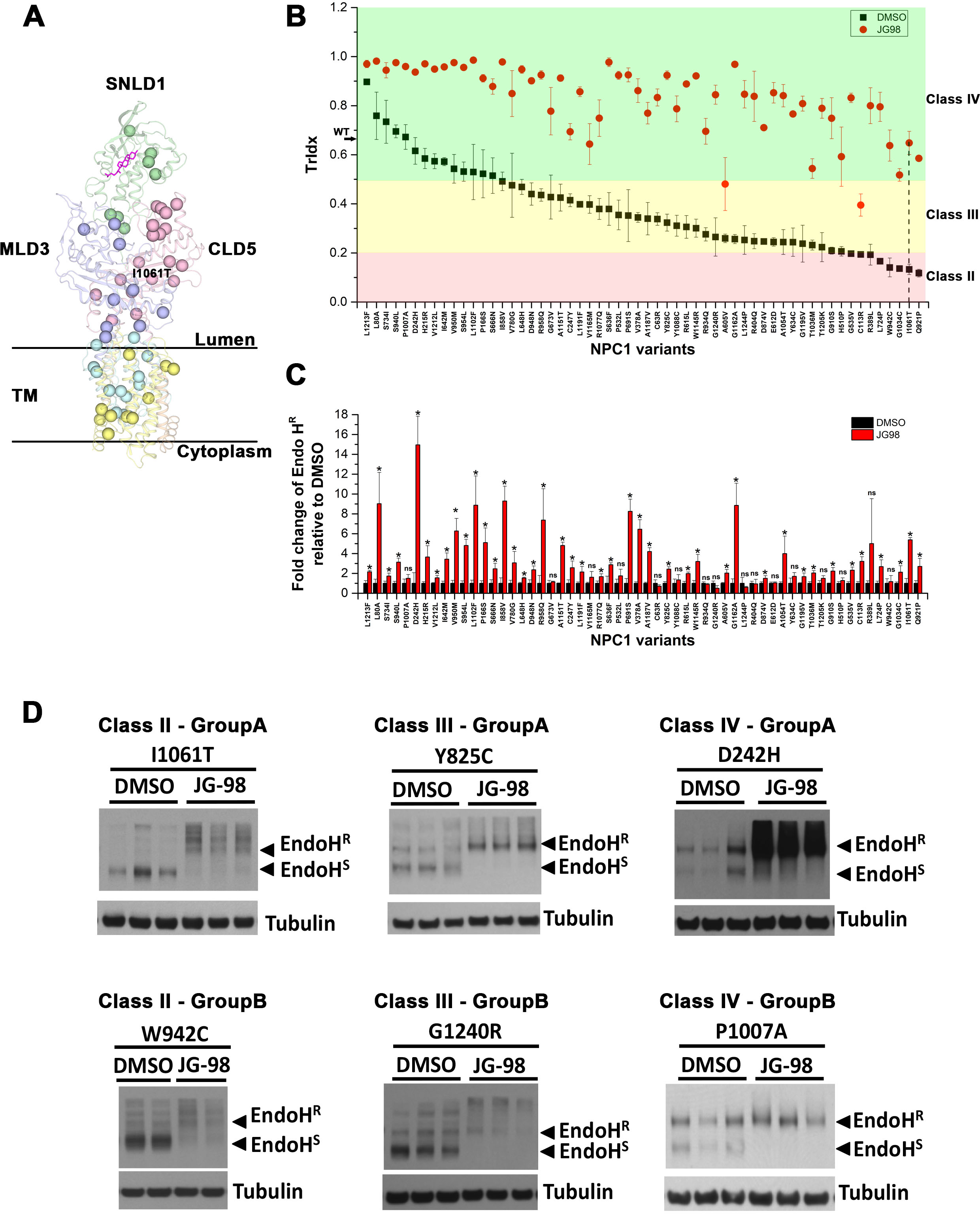
Impact of JG98 on the trafficking and stability of 58 NPC1 variants expressed in U2OS cells. (**A**) 58 NPC1 variants used in this study are showed as balls in NPC1 structure [13–15]. Each domain is labeled. Cholesterol is shown as a stick figure in magenta. (**B**) Quantification of the TrIdx for all the 58 variants in the absence (black squares) or presence (red circles) of JG98. Error bar is indicated as mean ± SD. Plasmid harboring each of the 58 variants was transfected in U2OS-SRA-shNPC1 cells. After treatment of 5 μM JG98 for 24 h, all sample lysates were treated with EndoH followed by Western blotting. Variants are ordered according to their TrIdx in the DMSO condition. Variants are grouped into different classes based on their trafficking index as indicated by background color (pink—Class II, yellow—Class III, green—Class IV). The p-value associated with each variant when compared to the JG98 treated or untreated condition was <0.05 (Student’s t-test). (**C**) Quantification of the fold-change of EndoH^R^ relative to DMSO after treatment of JG98 (mean ± SD). Variants are in the same order as (B). p-value for each variant compared to JG98 treated and untreated groups are labeled (p<0.05,*; p>0.05, ns; Student’s t-test). (**D**) Examples of Immunoblots for variants with increased EndoH^R^ (Group A) or no significant change of EndoH^R^ (Group B) in each TrIdx class are shown.

An analysis of the steady-state TrIdx allowed us to categorize NPC1 variants into one of four classes (Table 1). Class I includes a single variant that did not produce any NPC1 protein, representative of variants resulting in large protein truncations. Class II represents variants with a severe defect in trafficking efficiency (TrIdx<0.2), including I1061T variant (Fig3B, vertical dash line). Class III represents variants with intermediate trafficking (0.2<TrIdx<0.5) (Fig 3B). Class IV includes variants with > 50% of trafficking out of ER, including NPC1-WT that has ~0.65 TrIdx when expressed in U2OS cells (Fig. 3B, black arrow on y-axis). Strikingly, JG98 significantly improves the TrIdx of all the variants (Fig. 3B, red circles) reaching or exceeding WT-level, indicating that the Hsp70 system can play a prominent role in managing the stability and trafficking of NPC1 variants distributed across the NPC1 polypeptide.

**Table 1.** Classification of NPC1 variants according to their TrIdx when transiently transfected in the U2OS-SRA-shNPC1 cell line.

| **Class II: <20% R band** | **Class III: 20-50% R band** | **Class IV: >50% R band** |
| --- | --- | --- |
| C113R<br>R389L<br>G535V<br>L724P<br>Q921P<br>W942C<br>G1034C<br>I1061T | C63R<br>C247Y<br>V378A<br>R404Q<br>H510P<br>P532L<br>A605V<br>E612D<br>R615L<br>Y634C<br>S636F<br>L648H<br>G673V<br>P691S<br>V780G<br>Y825C<br>I858V<br>D874V<br>G910S<br>R934Q<br>D948N<br>R958Q<br>T1036M<br>A1054T<br>R1077Q<br>Y1088C<br>W1145R<br>A1151T<br>G1162A<br>V1165M<br>A1187V<br>L1191F<br>G1195V<br>T1205K<br>G1240R<br>L1244P | L80V<br>P166S<br>H215R<br>D242H<br>S666N<br>I642M<br>L748H<br>S734I<br>I858V<br>S940L<br>V950M<br>S954L<br>R1077Q<br>G1195V<br>L1213F<br>P1007A<br>T1205K |

Interestingly, though the trafficking efficiency (TrIdx) of all the variants expressed in U2OS cells is improved by JG98, there is a heterogeneity of responses regarding the level of EndoH^R^ glycoform (Fig 3C). The EndoH^R^ glycoforms are significantly increased for 42 variants while the changes of EndoH^R^ are not significant for 16 of the variants, despite concurrent improvement in TrIdx (Fig 3C). We further classified those variants that were significantly changed as Group A, while those that were not significantly changed were binned as Group B (Table 2). For example, as shown in Figure 3D, I1061T, Y825C and D242H (Class A) show a significant increase of EndoH^R^ band and decrease of Endo H^S^ band, conversely, W942C, G1240R and P1007A (Class B) only show significant clearance of Endo H^S^ yet without any significant changes on EndoH^R^ band. Taken together, these data suggest that the Hsp70 chaperone system differentially impacts the trafficking efficiency (TrIdx) and post-ER (EndoH^R^) stability of NPC1 variants on a residue-by-residue basis.

**Table 2.** Classification of NPC1 variants according to the response of the EndoH^R^ glycoform to JG98. Group A represents the variants with EndoH^R^ species that are significantly increased by JG98 treatment, while Group B represents the variants where the EndoH^R^ species are not changed significantly.

| Class II |  | Class III |  | Class IV |  |
| --- | --- | --- | --- | --- | --- |
| <u>Group A</u> | <u>Group B</u> | <u>Group A</u> | <u>Group B</u> | <u>Group A</u> | <u>Group B</u> |
| C113R | R389L | C247Y | C63R | L80A | P1007A |
| G535V | W942C | V378A | R404Q | P166S |  |
| L724P |  | A605V | H510P | H215R |  |
| Q921P |  | R615L | P532L | D242H |  |
| G1034C |  | S636F | E612D | I642M |  |
| I1061T |  | L648H | Y634C | S666N |  |
|  |  | P691S | G673V | S734I |  |
|  |  | V780G | R934Q | S940L |  |
|  |  | Y825C | Y1088C | V950M |  |
|  |  | I858V | V1165M | S954L |  |
|  |  | D874V | T1205K | L1102F |  |
|  |  | G910S | G1240R | V1212L |  |
|  |  | D948N | L1244P | L1213F |  |
|  |  | R958Q |  |  |  |
|  |  | T1036M |  |  |  |
|  |  | A1054T |  |  |  |
|  |  | R1077Q |  |  |  |
|  |  | W1145R |  |  |  |
|  |  | A1151T |  |  |  |
|  |  | G1162A |  |  |  |
|  |  | A1187V |  |  |  |
|  |  | L1191F |  |  |  |
|  |  | G1195V |  |  |  |

### Mapping JG98 response across entire NPC1 polypeptide through VSP

To provide a more in depth understanding of how the Hsp70 system manages the trafficking and post-ER stability of the entire NPC1 polypeptide and how these different functional features correlate with the NPC1 structure, likely reflecting proteostasis sensitive modulation of the protein fold, we applied variation spatial profiling (VSP) [2] to map the response of each residue to JG98 across the entire NPC1 polypeptide. This approach is based on the input value of the 58 variants that were experimentally measured (Fig 3). Briefly, VSP is a GP-based, machine learning (ML) approach that can utilize a sparse collection of variants to generate sequence-to-function-to-structure relationships at atomic resolution [2]. VSP is based on the new biological principle of spatial covariance (SCV) [2], which assigns set-points to each residue in the polypeptide chain to determine the role of that residue in sequence-to-function-to-structure design. In this approach, a matrix relationship is built based on SCV relationships of the experimentally measured sparse collection of variants to predict the unknown functional role(s) of amino acids spanning the rest of the NPC1 polypeptide chain. VSP generates phenotype landscapes that quantitatively assign the known and unknown SCV tolerance of each residue in the NPC1 fold to cellular physiology [2] to, in this case, interrogate directly their responses to JG98. These values can be directly mapped to the NPC1 structure [2] to provide a facile interpretation of the SCV relationships responsible for NPC1 trafficking.

Here, we used VSP to weight the SCV relationships according to their sequence position on the primary NPC1 sequence (x-axis) and plotted them against their relative, calculated TrIdx in the DMSO condition (y-axis) to build a phenotype landscape for delta (Δ) TrIdx in response to JG98 (z-axis) (Fig 4A, lower panel, r=0.4, p=0.002). Similarly, we assembled a map depicting the log_2_ fold change of EndoH^R^ band in response to JG98 (z-axis) (Fig 4D, lower panel, r=0.3, p=0.02). The highest confidence value for each residue in the phenotype landscape was mapped onto the NPC1 3D structure (Fig 4B-C and **4E-F**) [13, 14] to allow more intuitive description of NPC1 trafficking as a ‘functional-structure’ [2].

**Figure 4.**
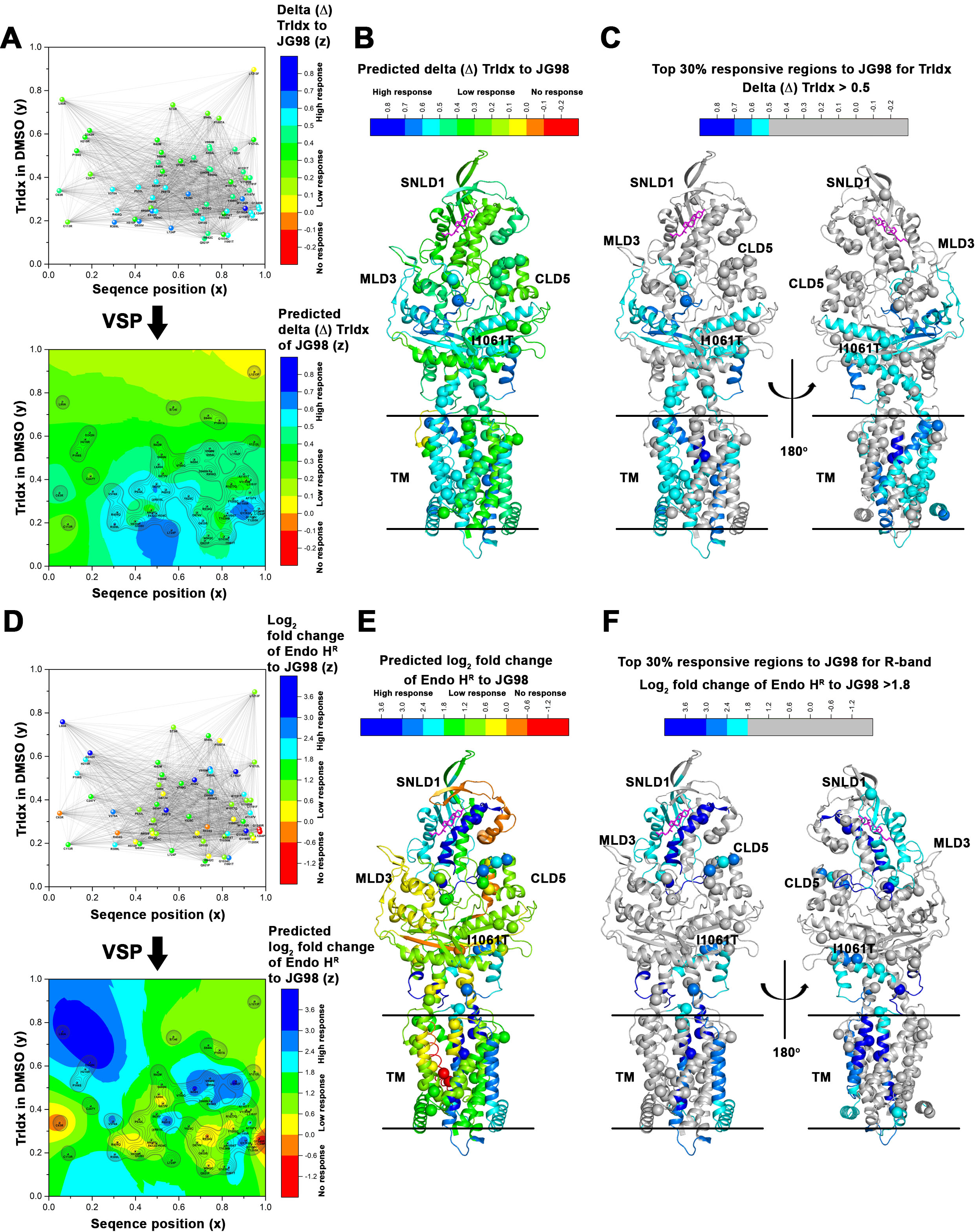
Mapping JG98 impact across the entire NPC1 polypeptide using VSP. (**A**) By analyzing the spatial covariance (SCV) relationships of 58 NPC1 variants according to their position in primary sequence (x-axis), TrIdx value in DMSO (y-axis) and delta (Δ) TrIdx response to JG98 (upper panel), VSP [2] constructs a phenotype landscape (lower panel) that maps delta (Δ) TrIdx response for all the residues across the entire NPC1 polypeptide. Color scale shows the predicted delta (Δ) TrIdx values with red-orange representing no response, yellow-green representing a low to medium response and blue-cyan representing a high response. Contour lines represent the confidence for the delta (Δ) TrIdx response prediction. (**B**) The phenotype landscape can be mapped to a structure snapshot of NPC1 [13–15] to show the specific structural features of NPC1 fold that JG98 targets. (**C**) The top 30% predicted TrIdx responsive regions to JG98 are highlighted by assigning the color for all regions below the 30% threshold TrIdx as gray. (**D-F**) VSP predicts the phenotype landscape for EndoH^R^ changes in response to JG98 (**D**). Log_2_ transformation of the fold change of EndoH^R^ after JG98 treatment is used as input for VSP training (upper panel) to generate the phenotype landscape (lower panel). The phenotype landscape is mapped onto structure (**E**). (**F**) The top 30% predicted EndoH^R^ responsive regions to JG98 are highlighted by assigning the color for all regions below the 30% threshold of EndoH^R^ response as gray.

Examination of these results showed that the delta (Δ) TrIdx landscape predicts that the TrIdx of almost all of the variants will be improved by JG98 (Fig 4A, lower panel, green-cyan-blue). Looking at the structure (Fig 4C), the top 30% high responsive regions (Fig 4A, lower panel, cyan-blue) generally have low TrIdx values in the DMSO condition (Fig 4A, lower panel, y=~0.2) and are principally localized to the transmembrane (TM) region (Fig 1A; Fig 4C) and the interface between MLD3 and CLD5 domains (Fig 1A; Fig 4C). The SCV relationships of these variant residues create ER SCV intolerance and thus lead to a severe trafficking deficiency. Strikingly, JG98 is able to increase the ER SCV tolerance to rescue their trafficking efficiency.

In contrast, the EndoH^R^ landscape was more complex, with both non-responsive regions (Fig 4D, lower panel, red-orange-yellow) and highly responsive regions (Fig 4D, lower panel, cyan-blue; Fig 4F). Most of the highly responsive regions have high TrIdx values in DMSO (Fig 4D, lower panel, cyan-blue, y=~0.6). For example, these regions are found in the SNLD1 domain (Fig 1A) that form the cholesterol-binding pocket (Fig 4F, cholesterol highlighted as magenta). Interestingly, variants at these residues have trafficking levels equivalent or better then WT. Thus, the dramatic increase in EndoH^R^ glycoforms in response to JG98 may indicate that the Hsp70 system normally plays a role in regulating the turnover rate of these NPC1 variants in the LE/Ly endomembrane system. Most of the residues that have low TrIdx values in the DMSO condition do not show a high response for EndoH^R^ glycoforms to JG98 (Fig 4D, lower panel). One possible explanation is that, although the TrIdx of those variants is improved (Fig 4A, lower panel), they are still unstable and JG98 fails to reverse the pathways leading to their degradation. One of the exceptions is the cluster contributed by A1054T, I1061T and G1162A, which links CLD5 (Fig 1A) to the TM region (Fig 1A; Fig. 4F), indicating that these particular mutants are potential strong candidates to be rescued by JG98. Together, these studies based on the computational formalism of VSP to link sequence-to-function-to-structure relationships [2], suggest that Hsp70 dynamically manages the proteostatic response of different NPC1 domains on a residue-by-residue basis.

### Impact of Hsp70 co-chaperone BAG on NPC1-I1061T trafficking

Because JG98 is an allosteric inhibitor that disrupts the protein-protein interaction between Hsp70 and BAG proteins, we silenced the major BAG family isoforms (BAG1-3) to test their effects on NPC1-I1061T (Fig 5A). This experiment is important as a genetic test of the selectivity of JG98 in this system. Similar to what we found with JG98, silencing of BAG1-3 either had little effect on TrIdx (siBAG3) or no effect (siBAG1-2) (Fig 5B, left panel). Thus, the Hsp70-BAG complex likely has a negligible impact on whether NPC1 continues through the export pathway or is selected for degradation. In contrast, the EndoH^R^ species of NPC1-I1061T are substantially (~3-fold) increased by silencing each of BAG1-3 (Fig 5B, right panel), indicating that BAG1-3 play a role in destabilizing NPC1-I1061T. Consistent with this observation, overexpression of BAG1 and BAG3 decrease the overall stability of NPC1-I1061T (Fig 5D; Fig 5E, right panel). These observations are consistent with the roles of BAG1 and BAG3 in mediating degradation of Hsp70 clients through the ubiquitin proteasome system (UPS) or autophagy-lysosomal pathway, respectively [37–39, 60–62]. Interesting, while the overall stability of NPC1-I1061T is reduced, the trafficking efficiency (TrIdx) is increased (Fig 5D, long exposure; Fig 5E, left panel), indicating there may be a balance in managing TrIdx and the protein level of NPC1 variants by BAGs. The impact of BAG1 and 3 on NPC1-I1061T is dependent on the BAG domain, as overexpression of ΔC44 of BAG1 and ΔC450 of BAG3 with BAG domain truncated (Fig 5C) stabilize NPC1-I1061T (Fig 5D; Fig 5E, left panel). Overexpression of BAG2 stabilized NPC1-I1061T, an observation that may relate to its inhibitory role on CHIP to facilitate the chaperone function of Hsp70. Together, these results indicate that BAG co-chaperones manage the overall SCV tolerance of the fold for degradation and contribute to the mechanism of JG98 impact.

**Figure 5.**
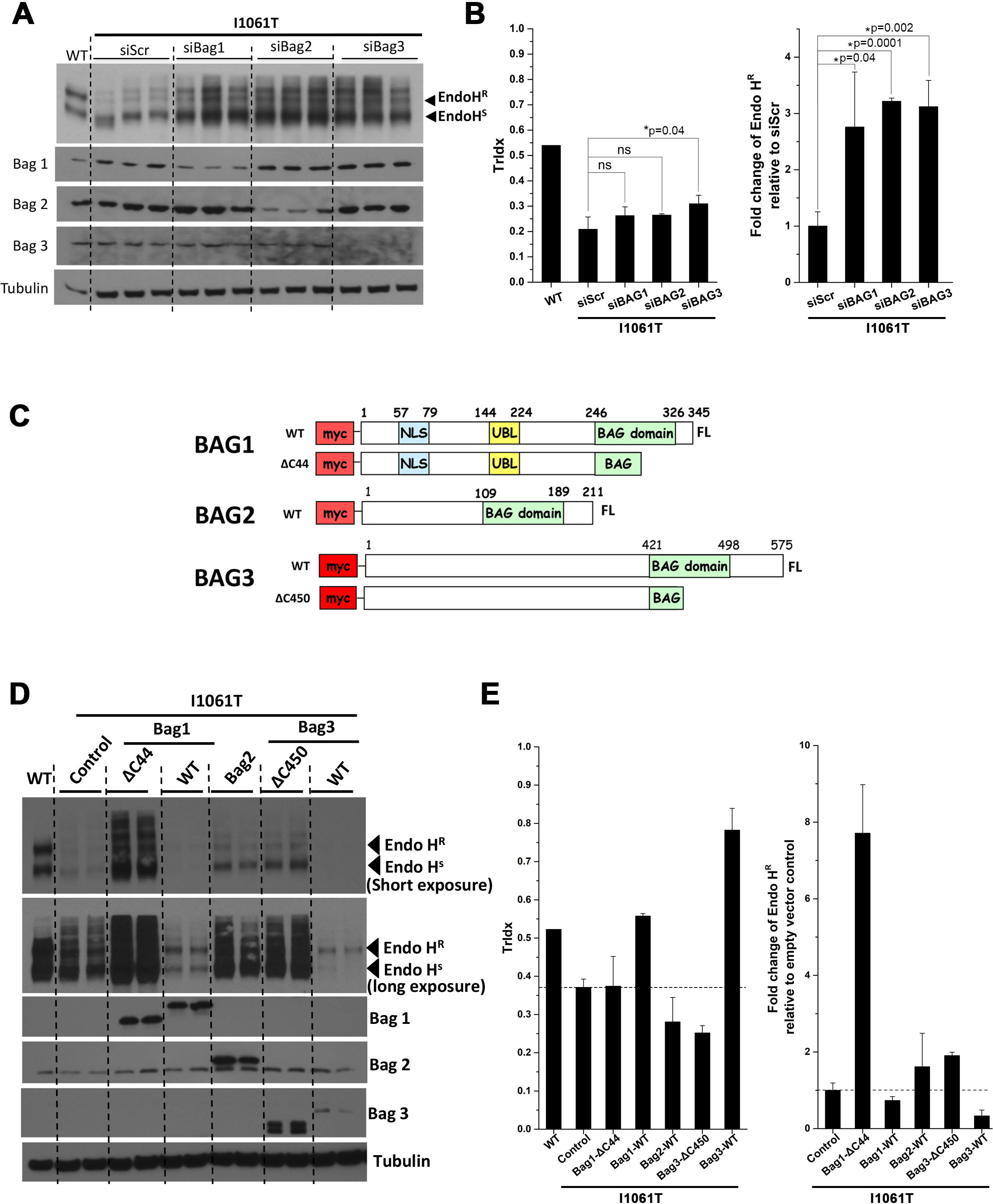
Impact of Hsp70 co-chaperone BAG proteins on NPC1-I1061T. (**A**) Immunoblot analysis EndoH digestion products of NPC1-I1061T after partial silencing of BAGs 1-3 with siRNA. (**B**) Quantification of (**A**) for TrIdx (left panel) and fold-change of EndoH^R^ relative to scramble siRNA control (right panel). The error bar (mean ± SD) and p-value (p<0.05, *; p>0.05, ns; Student’s t-test) are indicated. (**C**) Cartoon description of WT and modified BAG1-3 plasmids used for overexpression analysis. BAG1 has a deletion at ΔC44 which creates a truncated form of the protein with the BAG domain disrupted. Similarly, BAG3 has a deletion at ΔC450 which creates a BAG domain truncated form of the protein. (**D-E**) Immunoblot analysis (**D**) and quantitation (**E**) of EndoH digested products of NPC1-I1061T in response to overexpression of BAG proteins described in (**C**).

## DISCUSSION

We have provided the first description of the corrective properties of the Hsp70 allosteric inhibitor, JG98, on the trafficking of disease-associated variants of NPC1. At steady state, NPC1 variants fall into 1 of 4 categories based on their level of trafficking, including the class II (no trafficking), class III (moderate trafficking) and class IV (WT-like trafficking). Disease-causing class IV mutations likely represent variants with reduced or no activity(s) in the downstream endomembrane trafficking pathways and/or cholesterol management in the LE/Ly. Each of the variant populations were found to significantly, but differentially, respond to JG98. While the observations of improved TrIdx and EndoH^R^ levels following JG98 treatment are generally consistent with our observations in patient-derived fibroblasts, we did observe some inconsistencies, particularly for TrIdx. One possibility could stem from the 2-3-fold lower NPC1 expression seen in primary fibroblasts relative to that seen in transiently transfected U2OS cells, which required immunoprecipitation of NPC1, thereby possibly imposing a bias for the detection of EndoH^S^ versus EndoH^R^ species. A second possibility is that the primary fibroblast models are more restricted in their JG98-mediated proteostatic reprogramming. A third possibility is that each patient fibroblast carries the burden of its unique genomic diversity that could differentially impact how JG98 impacts NPC1 function; for example, by expressing different drug transporters. There could also be different off-target effects at higher concentration in each fibroblast line. Together, these results suggest that Hsp70 chaperone/co-chaperone system may target multiple pathways and that the impact of JG98 for NPC1 variants in different patients will not only be dependent on the patient specific variant but also could be affected by cell and tissue specific modifiers.

To address the complication of complexity in JG98 responses, we applied the computational formalism afforded by VSP that embraces diversity to explain biology [2]. This approach allows us to define the sequence-to-function-to-structure relationships through the new biological principle of SCV [2]. SCV is a universal metric that allows us to use a sparse collection of known variants and their functional attributes to predict the role of each amino acid in the NPC1 polypeptide through the matrix-based generation of phenotype landscapes. VSP is a powerful tool to deal with diversity in the population. Differences reflected in the dynamics of JG98-mediated fold management likely encompass the chemical, biochemical and biophysical features driving assembly and differential stability of the NPC1 polypeptide fold in the different compartments of the endomembrane system. Moreover, each variant expressing cell may harbor distinct activation states of their unfolded protein response (UPR) [63, 64] and heat-shock response (HSR) [26] pathways given that each variant likely presents a distinct folding problem leading to unique tuning of the maladaptive stress response (MSR) [24] that could further impact the response to JG98. Despite these complexities, the dynamic temporal and spatial relationships that dictate the SCV tolerance of the fold for a given environment reveal the impact of a variant on its functional-structure [2], each residue of which can be assigned a quantitative SCV set-point value in the phenotype landscape. Our VSP approach suggests that targeting specific protein-protein interaction between Hsp70 and its co-chaperones to adjust the SCV tolerance threshold for the residue of interest will reveal how to bin human variation from a more global, functional perspective.

A challenge of our studies is to understand how a cytosolic chaperone system impacts trafficking of a largely luminal protein such as NPC1. One possibility is that JG98 shifts the functional equilibrium between Hsp70 and the cytosolic ERAD or autophagy-lysosomal degradation pathways favoring productive chaperoning. A second possibility is that the JG98-mediated alterations in cytosolic Hsp70 chaperone activity triggers stress response pathways, including the UPR response, that will alter the level of ER luminal chaperone components that impact NPC1 variant folding. A third possibility is that JG98 is affecting the activity of a yet-to-be identified NPC1 ‘receptor’ for delivery to the COPII ER export machinery [65]. A fourth possibility is that JG98 is altering the steady state plasticity and selectively of endomembrane trafficking compartments through the activity of Rab GTPases, tether assemblies and SNARE complexes that manage compartmental design [65–67]. While how each of these possibilities contributes to the JG98-mediated correction of NPC1 variants remains unknown, they illustrate the complexity of the proteostasis buffering capacity in a given cell type that can be quantitated by a formalism that embraces diversity [2]. Quantitative SCV afforded by VSP mapping suggests that the Hsp70 chaperone system is a very dynamic process that tracks the assembly and maintenance of the polypeptide chain on a residue-by-residue basis-working as a continuous ‘*touch-n-go*’ machine-looking at each residue in the context of their local kinetic and/or thermodynamic features, a conclusion supported by the known dynamic structural features of the Hsp70 cycle [29, 34, 52].

In summary, VSP [2] links sequence-to-function-to-structure relationships to understand the genotype to phenotype transformation occurring in response to JG98. These results indicate that targeting specific protein-protein interactions between Hsp70 and its co-chaperones may have broad therapeutic potential for NPC disease, albeit one that will need to be handicapped by knowledge in advance of the sensitivity of each residue in the polypeptide chain to a therapeutic, insights which can be revealed by VSP-based analyses. We have posited that this new way of thinking alters our perspective of central dogma from a linear (DNA<->RNA->Protein) to that of a matrix (SCV[DNA<->RNA->Protein] way of thinking [2], one which will likely require more advanced GP-based generative modeling to precision map the impact of a therapeutic in misfolding disease in the individual [2].

## MATERIALS AND METHODS

### Reagents

Small molecule Hsp70 modulators (JG compounds) were a generous gift from Jason Gestwicki (San Francisco, CA). Endoglycosidase H (EndoHf) was purchased from New England Biolabs (Ipswich, MA). The primary antibodies used were rat monoclonal hNPC-1 created in-house and rat monoclonal anti-tubulin antibody from Abcam (Cambridge, MA). Benzonase Nuclease and cOmplete-Mini Protease Inhibitor Cocktail was used for the cell lysis and purchased from Sigma (St. Louis, MO).

### Cell lines and cell culture

Human Wild-Type (GM05659), homozygous NPC1^I1061T^ (GM18453), heterozygous fibroblasts NPC1^T1036M/P1007A^ (GM17912), NPC1^G673V/I1061T^ (GM18398), NPC1^V1165M/I1061T^ (GM18398), NPC1^P401T/I1061T^ (GM17920), NPC1^N1165S/R1186H^ (GM18397) were purchased from Coriell Cell Repositories (Coriell Institute of Medical Research). Additionally, the following heterozygous fibroblasts NPC1^I1061T/R1186G^ (TQNPC5), NPC1^I1061T/M1142T^ (ARNPC9), NPC1^1036M/S954L^ (CRNPC67), NPC1^R404Q/P1007A^ (TWRNPC60), NPC1^I1061T/D948N^ (MONPC63), NPC1^I1061T/P1007A^ (MBNPC44), NPC1^I1061T/R404Q^ (TLNPC36), NPC1^I1061T/I1094T^ (TBNPC40), NPC1^I1061T/S954L^ (KWNPC53), NPC1^C177Y/V950M^ (JSNPC49) were courtesy of Denny Porter Lab (NIH, Maryland) The cells were grown in DMEM medium with 2 mM L-glutamine, 10% FBS, 50 units/ml penicillin, and 50 μg/ml streptomycin antibiotics. To examine a large number of NPC1 variants, the cell line U2OS-SRA-shNPC1 was used, in which NPC1 expression is stably silenced [20] The antibiotic puromycin was used for selection. Stable lines were grown in McCoy’s 5A medium (1X: modified) with 5% FBS, 3μg puromycin and 1mg/ml G418.

### NPC1 transfection in U2OS-SRA-shNPC1

Cells were seeded at 2.0 ×10^5^ cells/ml in 12-well plates and incubated overnight. In order to express NPC1, transfections was performed using the reagent FuGENE 6 from PROMEGA (Madison, WI) with a 1:4 ratio, 2μg DNA: 8μL FuGENE6 in Opti-MEM (1X) + Hepes + Sodium Bicarbonate + L-glutamine + 5% FBS. Samples were then incubated for 5 h at 37°C and the medium was replaced with normal growth medium. Cells were then incubated for another 48 h and visualized under the fluorescent microscope for GFP-positive transfected cells. These variants were all transiently expressed in the U2OS-SRA-shNPC1 cells. After 72 h from initial plating, cells were dosed with either dimethyl sulfoxide (DMSO) as small vehicle control, or small molecule JG series Hsp70 allosteric modulators in normal growth media at 5 μM and incubated for 24 h at 37°C. After a total of 96 h, cells washed twice with 1X PBS and stored at −80°C until lysis. In order to collect statistically significant data results, each variant and drug treatment condition were performed in triplicate. Additionally, to examine overexpression of Bag1 and 2 and their effect on NPC1 protein trafficking, a co-transfection was done following the same protocol as seen above, but with 1 μg of NPC1 plasmid + 1μg of Bag1-3 plasmids.

### Transfection of NPC1 variants and silencing of BAGs

Cells were seeded at 2.0 × 10^5^ cells/ml in 12-well plates and incubated overnight. In order to silence the Bag1-3, Lipofectamine RNAi MAX (Thermofisher) was used to deliver the siRNA. The RNAi MAX and siRNA were diluted in Opti-MEM (1X) + Hepes + sodium bicarbonate + L-glutamine + 5% FBS at 1:100 and 1:50 respectively. Samples were incubated at rt for 20 min and then added to cells and incubated at 37°C for 5 h. Medium was then changed to normal growth medium and incubated for 24 h. Transfection of NPC1 variants was then conducted (a total of 48 h after being seeded) using FuGENE 6 from PROMEGA (Madison, WI). Ratio was 1:4, 2μg DNA: 8uL FuGENE6 in Opti-MEM (1X) + Hepes + sodium bicarbonate + L-glutamine + 5% FBS. The 12-well plate was incubated for 5 h and then changed to normal growth medium. Cells were then incubated for another 48 h and visualized under the fluorescent microscope for transfected cells (GFP) every 24 h. Cells were washed twice with 1X PBS and stored at −80°C until lysis.

### Sample preparation

Cells were lysed with 50 μL/well of 1X RIPA (150 mM NaCl, 1.0% IGEPAL Ca-360, 0.5%Na-deoxychlorate, O.1% SDS, 50 mM Tris pH 8.0, 1X protease inhibitors, 1X benzonase) on ice for 30 min. Samples were collected and incubated at 37°C for 20 min, centrifuged at 14K RPM at 4°C for 20 min. Supernatant was collected and the BCA assay performed. EndoH digestion was performed by addition of 1X glycoprotein denaturing buffer (New England Biolabs) to 20 μg of lysate and incubated at 50°C for 30 min. For each sample 10,000 U/ml EndoHf and 1X Glycobuffer (New England Biolabs) was added and incubated overnight at 37°C. Protein samples were separated on 4-12% Bis-Tris BOLT protein gel (Invitrogen) and transferred to nitrocellulose membrane. Membranes were probed with NPC1 1:1000 and tubulin 1:25,000 diluted in 5% milk solution.

For patient fibroblast samples, fibroblasts were seeded in a 6-well plate and grown until confluent. Once confluent, cells were treated with DMSO or JG-98 at 5 μM for 24 h at 37°C in normal growth media. Cells were then stopped and washed twice with 1X PBS and stored at −80°C until lysis. Following treatment with JG98 or vehicle cells were lysed with 125 μL/well of 1X RIPA (150 mM NaCl, 1.0% IGEPAL Ca-360, 0.5%Na-deoxychlorate, O.1% SDS, 50mM Tris pH 8.0, 1X Protease inhibitors, 1X Benzonase) on ice for 30 min. Samples were collected and then incubated at 37°C for 20 min. Samples were then centrifuged at 14K RPM at 4°C for 20 min. Supernatant was collected and BCA assay was performed. Based on the BCA Assay, 75 μg of each lysate sample was used for immunoprecipitation with a NPC1 specific antibody. Samples were incubated overnight with 15 μL of NPC1 Ab beads at 4°C with rotation. Lysate samples with beads were centrifuged at 500 RPM at 4°C for 3 min. Supernatant was removed and washed at 10-20X bead volume 2X (50 mM Tris-HCl, pH7.4; 150 mM NaCl; 1% Triton 100-X). After each wash samples were centrifuged at 500 RPM at 4°C for 3 min and the supernatant discarded. A final wash was performed at 10-20X bead volume 2X (50mM Tris-HCl, pH7.4; 150 mM NaCl). Lysate with beads was centrifuged at 500 RPM at 4°C for 3 min and the supernatant discarded. Samples were denatured with 35 μL of 1X Glycoprotein Denaturing buffer (New England Biolabs) at 95°C for 10 min. Denatured samples were centrifuged at 500 RPM at rt for 5 min. Supernatant was collected and 12 μL of sample was prepared for EndoH treatment (New England Biolabs). Samples were incubated with 10,000 U/ml EndoHf and 1X Glycobuffer (New England Biolabs) overnight at 37°C. Protein samples were separated on 4-12% Bis-Tris BOLT protein gel (Invitrogen) and transferred to nitrocellulose membrane. Membranes were probed with anti-NPC1 antibody at 1:1000 and anti-tubulin antibody at 1:25,000 diluted in 5% milk solution.

### NPC1 expression Vector

The cDNA encoding human ΔU3h*NPC1*-WT construct was kindly provided by Dan Ory (Washington University, St. Louis, MO). The NPC1-WT gene was cloned into the pMIEG3 vector, a murine stem cell virus (MSCV) retrovirus construct that allows the co-expression of GFP and a second gene, in this case NPC1. pMIEG3 plasmid was initially generated from pMSCVneo vector (Clontech) where the neomycin resistance site (Neo^r^) and the murine phosphoglycerate kinase promoter (P_PKG_), which controls expression of the neomycin resistance for antibiotic selection in eukaryotic cells, was removed. In its place the internal ribosome entry site (IRES) with enhanced green fluorescent protein (eGFP) gene was generated. The WT construct used in this study had four additional substitution when compared to the current reference NPC1 sequence. The mutations include 387 T>C (Y129Y), 1415 T>C (L472P), 1925 T>C (M642T), and 258 T>C (S863P). These variants were present in the original ΔU3h*NPC1*-WT construct, which has been previously used in the past as the WT protein control [42, 55–59]. This pMIEG3-hNPC1 plasmid was used as a template to generate all other NPC1 variants using the Quick-Change XL Site-directed Mutagenesis Kit (Stratagene, La Jolla, CA).

### Drug treatment of NPC1

Cells were incubated with DMSO (control) or small molecule Hsp70 modulators at 5 μM final concentration and incubated at 37° C for 24 h unless otherwise indicated.

### VSP analysis

Methods as described in [2].

## Acknowledgements

Support provided by the National Institutes of Health Grants HL095524 and DK051870 to WEB and NIR R01NS059690 and the Tau Consortium to JEG. We thank the continuing support of the Ara Parseghian Medical Research Foundation (APMRF) and Support for Accelerated Research (SOARs) for Samantha Scott and Dr. Pei Zhang.

## References Cited

1. Hindorff, L.A., et al., Prioritizing diversity in human genomics research. Nat Rev Genet, 2018. 19(3): p. 175–185.

2. Wang, C. and W.E., Balch, Bridging Genomics to Phenomics at Atomic Resolution through Variation Spatial Profiling. Cell Rep, 2018. 24(8): p. 2013–2028 e6.

3. Torkamani, A., et al., High-Definition Medicine. Cell, 2017. 170(5): p. 828–843.

4. Landrum, M.J., et al., ClinVar: improving access to variant interpretations and supporting evidence. Nucleic Acids Res, 2018. 46(D1): p. D1062–D1067.

5. Shao, H., et al., Exploration of Benzothiazole Rhodacyanines as Allosteric Inhibitors of Protein-Protein Interactions with Heat Shock Protein 70 (Hsp70). J Med Chem, 2018. 61(14): p. 6163–6177.

6. Moses, M.A., et al., Targeting the Hsp40/Hsp70 Chaperone Axis as a Novel Strategy to Treat Castration-Resistant Prostate Cancer. Cancer Res, 2018. 78(14): p. 4022–4035.

7. Balchin, D., M. Hayer-Hartl, and F.U., Hartl, In vivo aspects of protein folding and quality control. Science, 2016. 353(6294): p. aac4354.

8. Balch, W.E., et al., Adapting proteostasis for disease intervention. Science, 2008. 319(5865): p. 916–9.

9. Vance, J.E. and B. Karten, Niemann-Pick C disease and mobilization of lysosomal cholesterol by cyclodextrin. J Lipid Res, 2014. 55(8): p. 1609–21.

10. Vanier, M.T., Niemann-Pick diseases. Handb Clin Neurol, 2013. 113: p. 1717–21.

11. Vanier, M.T., Complex lipid trafficking in Niemann-Pick disease type C. J Inherit Metab Dis, 2015. 38(1): p. 187–99.

12. Vanier, M.T., Niemann-Pick disease type C. Orphanet J Rare Dis, 2010. 5: p. 16.

13. Li, X., et al., 3.3 A structure of Niemann-Pick C1 protein reveals insights into the function of the C-terminal luminal domain in cholesterol transport. Proc Natl Acad Sci U S A, 2017. 114(34): p. 9116–9121.

14. Li, X., et al., Structure of human Niemann-Pick C1 protein. Proc Natl Acad Sci U S A, 2016. 113(29): p. 8212–7.

15. Gong, X., et al., Structural Insights into the Niemann-Pick C1 (NPC1)-Mediated Cholesterol Transfer and Ebola Infection. Cell, 2016. 165(6): p. 1467–1478.

16. Gelsthorpe, M.E., et al., Niemann-Pick type C1I1061T mutant encodes a functional protein that is selected for endoplasmic reticulum-associated degradation due to protein misfolding. J Biol Chem, 2008. 283(13): p. 8229–36.

17. Yu, T., et al., Ryanodine receptor antagonists adapt NPC1 proteostasis to ameliorate lipid storage in Niemann-Pick type C disease fibroblasts. Hum Mol Genet, 2012. 21(14): p. 3205–14.

18. Ohgane, K., et al., Discovery of oxysterol-derived pharmacological chaperones for NPC1: implication for the existence of second sterol-binding site. Chem Biol, 2013. 20(3): p. 391–402.

19. Ebrahimi-Fakhari, D., et al., Reduction of TMEM97 increases NPC1 protein levels and restores cholesterol trafficking in Niemann-pick type C1 disease cells. Hum Mol Genet, 2016. 25(16): p. 3588–3599.

20. Pipalia, N.H., et al., Histone deacetylase inhibitors correct the cholesterol storage defect in most Niemann-Pick C1 mutant cells. J Lipid Res, 2017. 58(4): p. 695–708.

21. Calamini, B., et al., Small-molecule proteostasis regulators for protein conformational diseases. Nat Chem Biol, 2011. 8(2): p. 185–96.

22. Hutt, D.M., E.T., Powers, and W.E., Balch, The proteostasis boundary in misfolding diseases of membrane traffic. FEBS Lett, 2009. 583(16): p. 2639–46.

23. Rauniyar, N., et al., Quantitative Proteomics of Human Fibroblasts with I1061T Mutation in Niemann-Pick C1 (NPC1) Protein Provides Insights into the Disease Pathogenesis. Mol Cell Proteomics, 2015. 14(7): p. 1734–49.

24. Roth, D.M., et al., Modulation of the maladaptive stress response to manage diseases of protein folding. PLoS Biol, 2014. 12(11): p. e1001998.

25. Balch, W.E., D.M., Roth, and D.M., Hutt, Emergent properties of proteostasis in managing cystic fibrosis. Cold Spring Harb Perspect Biol, 2011. 3(2).

26. Li, J., J. Labbadia, and R.I., Morimoto, Rethinking HSF1 in Stress, Development, and Organismal Health. Trends Cell Biol, 2017. 27(12): p. 895–905.

27. Sala, A.J., L.C., Bott, and R.I., Morimoto, Shaping proteostasis at the cellular, tissue, and organismal level. J Cell Biol, 2017. 216(5): p. 1231–1241.

28. Labbadia, J. and R.I., Morimoto, The biology of proteostasis in aging and disease. Annu Rev Biochem, 2015. 84: p. 435–64.

29. Zuiderweg, E.R., L.E., Hightower, and J.E., Gestwicki, The remarkable multivalency of the Hsp70 chaperones. Cell Stress Chaperones, 2017. 22(2): p. 173–189.

30. Morozova, K., et al., Structural and Biological Interaction of hsc-70 Protein with Phosphatidylserine in Endosomal Microautophagy. J Biol Chem, 2016. 291(35): p. 18096–106.

31. Pratt, W.B., et al., Targeting Hsp90/Hsp70-based protein quality control for treatment of adult onset neurodegenerative diseases. Annu Rev Pharmacol Toxicol, 2015. 55: p. 353–71.

32. Zarouchlioti, C., et al., DNAJ Proteins in neurodegeneration: essential and protective factors. Philos Trans R Soc Lond B Biol Sci, 2018. 373(1738).

33. Gorenberg, E.L. and S.S., Chandra, The Role of Co-chaperones in Synaptic Proteostasis and Neurodegenerative Disease. Front Neurosci, 2017. 11: p. 248.

34. Craig, E.A. and J. Marszalek, How Do J-Proteins Get Hsp70 to Do So Many Different Things? Trends Biochem Sci, 2017. 42(5): p. 355–368.

35. Brehme, M. and C. Voisine, Model systems of protein-misfolding diseases reveal chaperone modifiers of proteotoxicity. Dis Model Mech, 2016. 9(8): p. 823–38.

36. Alderson, T.R., J.H., Kim, and J.L., Markley, Dynamical Structures of Hsp70 and Hsp70-Hsp40 Complexes. Structure, 2016. 24(7): p. 1014–30.

37. Rauch, J.N., E.R., Zuiderweg, and J.E., Gestwicki, Non-canonical Interactions between Heat Shock Cognate Protein 70 (Hsc70) and Bcl2-associated Anthanogene (BAG) Co-Chaperones Are Important for Client Release. J Biol Chem, 2016. 291(38): p. 19848–57.

38. Rauch, J.N. and J.E., Gestwicki, Binding of human nucleotide exchange factors to heat shock protein 70 (Hsp70) generates functionally distinct complexes in vitro. J Biol Chem, 2014. 289(3): p. 1402–14.

39. Hutt, D.M., et al., Silencing of the Hsp70-specific nucleotide-exchange factor BAG3 corrects the F508del-CFTR variant by restoring autophagy. J Biol Chem, 2018. 293(35): p. 13682–13695.

40. Kirkegaard, T., et al., Heat shock protein-based therapy as a potential candidate for treating the sphingolipidoses. Sci Transl Med, 2016. 8(355): p. 355ra118.

41. Schultz, M.L., K.L., Krus, and A.P., Lieberman, Lysosome and endoplasmic reticulum quality control pathways in Niemann-Pick type C disease. Brain Res, 2016. 1649(Pt B): p. 181–188.

42. Nakasone, N., et al., Endoplasmic reticulum-associated degradation of Niemann-Pick C1: evidence for the role of heat shock proteins and identification of lysine residues that accept ubiquitin. J Biol Chem, 2014. 289(28): p. 19714–25.

43. Petersen, N.H., et al., Connecting Hsp70, sphingolipid metabolism and lysosomal stability. Cell Cycle, 2010. 9(12): p. 2305–9.

44. Assimon, V.A., et al., Hsp70 protein complexes as drug targets. Curr Pharm Des, 2013. 19(3): p. 404–17.

45. Li, X., et al., Validation of the Hsp70-Bag3 protein-protein interaction as a potential therapeutic target in cancer. Mol Cancer Ther, 2015. 14(3): p. 642–8.

46. Li, X., et al., Analogs of the Allosteric Heat Shock Protein 70 (Hsp70) Inhibitor, MKT-077, as Anti-Cancer Agents. ACS Med Chem Lett, 2013. 4(11).

47. Wang, A.M., et al., Activation of Hsp70 reduces neurotoxicity by promoting polyglutamine protein degradation. Nat Chem Biol, 2013. 9(2): p. 112–8.

48. Taguwa, S., et al., Defining Hsp70 Subnetworks in Dengue Virus Replication Reveals Key Vulnerability in Flavivirus Infection. Cell, 2015. 163(5): p. 1108–1123.

49. Khachatoorian, R., et al., Allosteric heat shock protein 70 inhibitors block hepatitis C virus assembly. Int J Antimicrob Agents, 2016. 47(4): p. 289–96.

50. Young, Z.T., et al., Stabilizing the Hsp70-Tau Complex Promotes Turnover in Models of Tauopathy. Cell Chem Biol, 2016. 23(8): p. 992–1001.

51. Taguwa, S. and J. Frydman, The significance of Hsp70 subnetwork for Dengue virus lifecycle. Uirusu, 2015. 65(2): p. 179–186.

52. Zuiderweg, E.R., et al., Allostery in the Hsp70 chaperone proteins. Top Curr Chem, 2013. 328: p. 99–153.

53. Chung, C., et al., Genetic and pharmacological evidence implicates cathepsins in Niemann-Pick C cerebellar degeneration. Hum Mol Genet, 2016. 25(7): p. 1434–46.

54. Praggastis, M., et al., A murine Niemann-Pick C1 I1061T knock-in model recapitulates the pathological features of the most prevalent human disease allele. J Neurosci, 2015. 35(21): p. 8091–106.

55. Pipalia, N.H., et al., Histone deacetylase inhibitor treatment dramatically reduces cholesterol accumulation in Niemann-Pick type C1 mutant human fibroblasts. Proc Natl Acad Sci U S A, 2011. 108(14): p. 5620–5.

56. Millard, E.E., et al., Niemann-pick type C1 (NPC1) overexpression alters cellular cholesterol homeostasis. J Biol Chem, 2000. 275(49): p. 38445–51.

57. Zhang, M., et al., Cessation of rapid late endosomal tubulovesicular trafficking in Niemann-Pick type C1 disease. Proc Natl Acad Sci U S A, 2001. 98(8): p. 4466–71.

58. Watari, H., et al., Niemann-Pick C1 protein: obligatory roles for N-terminal domains and lysosomal targeting in cholesterol mobilization. Proc Natl Acad Sci U S A, 1999. 96(3): p. 805–10.

59. Watari, H., et al., Mutations in the leucine zipper motif and sterol-sensing domain inactivate the Niemann-Pick C1 glycoprotein. J Biol Chem, 1999. 274(31): p. 21861–6.

60. Li, F., et al., Exploring the multifaceted roles of heat shock protein B8 (HSPB8) in diseases. Eur J Cell Biol, 2018. 97(3): p. 216–229.

61. Liang, J., et al., Chaperone-Mediated Autophagy Protein BAG3 Negatively Regulates Ebola and Marburg VP40-Mediated Egress. PLoS Pathog, 2017. 13(1): p. e1006132.

62. Behl, C., Breaking BAG: The Co-Chaperone BAG3 in Health and Disease. Trends Pharmacol Sci, 2016. 37(8): p. 672–688.

63. Gardner, B.M., et al., Endoplasmic reticulum stress sensing in the unfolded protein response. Cold Spring Harb Perspect Biol, 2013. 5(3): p. a013169.

64. Plate, L. and R.L., Wiseman, Regulating Secretory Proteostasis through the Unfolded Protein Response: From Function to Therapy. Trends Cell Biol, 2017. 27(10): p. 722–737.

65. Aridor, M., COPII gets in shape: Lessons derived from morphological aspects of early secretion. Traffic, 2018.

66. Gurkan, C., A.V., Koulov, and W.E., Balch, An evolutionary perspective on eukaryotic membrane trafficking. Adv Exp Med Biol, 2007. 607: p. 73–83.

67. Gurkan, C., et al., Large-scale profiling of Rab GTPase trafficking networks: the membrome. Mol Biol Cell, 2005. 16(8): p. 3847–64.

